# Circadian gene *Bmal1* in the lateral habenula regulates alcohol drinking behavior in a sex specific manner

**DOI:** 10.1101/2024.08.13.607765

**Authors:** Cassandra Goldfarb, Vanessa Hasenhundl, Arianne Menasce, Shimon Amir, Konrad Schöttner

## Abstract

The circadian clock regulates most aspects of mammalian physiology and behaviour, including alcohol drinking behaviour. Disrupted circadian clock function via deletion of clock genes along the mesolimbic dopamine (DA) pathway has been linked to altered patterns of alcohol drinking behaviour. The lateral habenula (LHb), an epithalamic structure that house a semi-autonomous circadian clock, is a negative regulator of the mesolimbic DA system and of alcohol consumption. To study the role of the LHb circadian clock in alcohol consumption, we knocked out the core clock gene, *Bmal1* specifically in the LHb and examined the impact on various alcohol drinking paradigms and affective behaviours in male and female mice. Our findings demonstrate that *Bmal1* deletion in the LHb leads to sex-specific alterations in alcohol consumption. Male knockout mice exhibited increased voluntary alcohol intake, enhanced consumption of a bitter alcohol solution, and elevated alcohol binge drinking compared to controls. Conversely, female knockouts showed a marginal decrease in voluntary intake, significantly reduced consumption of an aversive alcohol solution, and lower post abstinence, relapse-like drinking. These results indicate that *Bmal1* in the LHb exerts a repressive effect on alcohol intake in males, while it facilitates intake under certain aversive conditions in females. Interestingly, *Bmal1* deletion did not significantly affect anxiety-like or depressive-like behaviours, suggesting that the habenular clock’s role in alcohol consumption is independent of affective state. These findings mark *Bmal1* in the LHb as a novel sexually dimorphic regulator of alcohol consumption in mice. Potential mechanisms involving the circadian modulation of DA and serotonin signaling pathways by the LHb clock is discussed.

## INTRODUCTION

Circadian rhythms are regulated by cellular clocks comprising a small set of so-called clock genes (Partch et al., 2014; Takahashi, 2017). These genes drive transcription and translation feedback loops that are at the core of the ∼24 rhythm generated by the clock. In mouse models, disruption or complete loss of function in different clock genes is associated with abnormal or total loss of circadian rhythmicity and in a range of physiological, metabolic, and behavioural disturbances (Bunger et al., 2000; King et al., 1997; Kume et al., 1999). In recent years, evidence has emerged linking clock genes, including Bmal1, Clock, Per2, and Rev-erba with alcohol drinking behaviour and alcohol use disorder (AUD). For example, polymorphisms in *Bmal1* and *Per2* have been linked to abnormal alcohol consumption, suggesting a clock gene based predisposition for developing AUD (Kovanen et al., 2010; Partonen, 2015; Valenzuela et al., 2016). In mice, constitutive knockouts of *Per2* and *Clock*, as well as selective suppression of *Clock* in the nucleus accumbens (NAc), can increase alcohol intake (Gamsby et al., 2013; Kirkpatrick et al., 2009; Rizk et al., 2022; Spanagel et al., 2005). Conversely, global knockout of *Rev-erba* decreases alcohol consumption in mice (Al-Sabagh et al., 2022), highlighting distinct roles for different clock genes in regulating alcohol drinking behaviour. More recently, we found that conditional deletion of *Bmal1* from medium spiny neurons (MSNs) throughout the striatum augments voluntary alcohol consumption and preference in male mice and conversely, repress intake and preference in females (de Zavalia et al., 2021). Interestingly, deletion of *Bmal1* only in the NAc augmented alcohol intake in both sexes (Herrera et al., 2023), indicating that *Bmal1* influences alcohol consumption via both shared, sex-independent mechanisms within the NAc and distinct, sex-specific mechanisms likely in the dorsal striatum.

It is unknown whether the sex dependent and independent effects of Bmal1 on alcohol consumption are specific to the striatum or whether they extend to other brain regions implicated in the control of alcohol drinking behaviour. In the present study we focused on the lateral habenula (LHb), a major negative regulator of the midbrain dopaminergic neurons that supply the striatum (Ji & Shepard, 2007; Matsumoto & Hikosaka, 2007; Salaberry & Mendoza, 2016), and a key player in aversive conditioning and alcohol drinking behaviour (Fu et al., 2017; Mondoloni et al., 2022; Salaberry & Mendoza, 2016; Zuo et al., 2014). Notably, the LHb houses a semi-autonomous circadian clock (Baño-otálora & Piggins, 2017; Guilding et al., 2010; Mendoza, 2017; Pradel et al., 2022; Sakhi et al., 2014), suggesting a potential influence on alcohol consumption via indirect circadian control of DA signaling in the striatum.

To study the role of the LHb clock in alcohol consumption, we investigated the effect of conditional deletion of *Bmal1* in the LHb on daily intermittent alcohol consumption, aversion-resistant consumption, reintroduction after forced abstinence, and alcohol binge drinking in male and female mice. Given the LHb’s involvement in affective behaviours (Hikosaka et al., 2008; Mendoza, 2017; Yang et al., 2018), depressive- and anxiety-like behaviours were assessed to identify potential interactions between *Bmal1*-mediated changes in alcohol consumption and affective state.

## RESULTS

### Knockout of *Bmal1* in the habenula

*Bmal1* floxed male and female mice received bilateral intra-LHb infusions of adeno associated virus (AAV) expressing Cre-recombinase and eGFP (KO) or AAV expressing eGFP only (CTR). Analysis of GFP and Cre expressing cells from histologically validated surgeries carried out at the end of the study revealed that ∼70% of cells in the LHb of experimental animals expressed Cre/eGFP (Fig. 1A). Immunofluorescence images confirm BMAL1 deletion in the habenula of KO mice (Fig. 1, B). Animals with low numbers of infected cells or off target infection were excluded from analyses.

**Figure 1.**
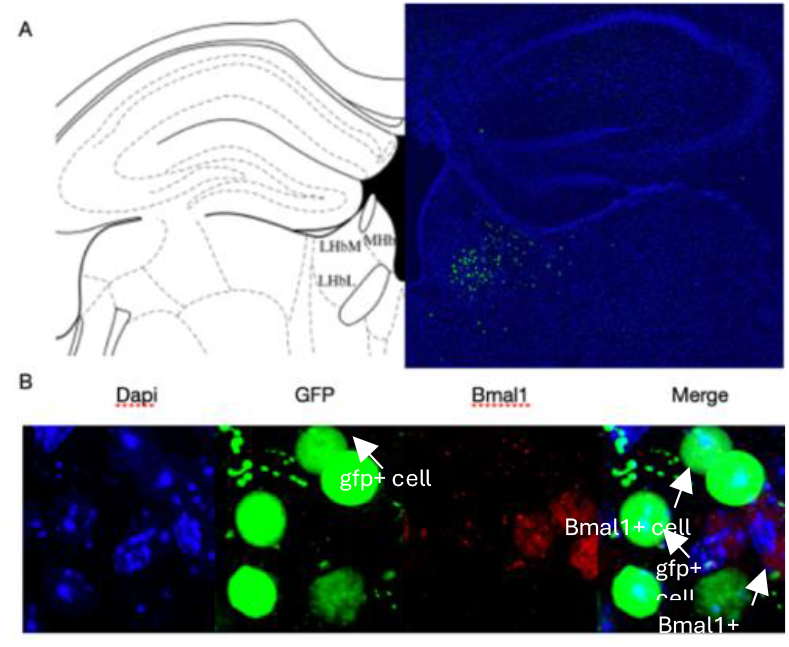
A) Representative immunofluorescence staining image of GFP expression in LHb of mice receiving stereotaxic injections of AAV-CAG-Cre-EGFP viral vector. GFP staining is centralized in the LHb. B) 60x representative image of Dapi (blue), GFP (green), and BMAL1 (red) immunofluorescence staining in LHb tissue of a knockout mouse. Arrows point to either GFP positive cell or a BMAL1 positive cell, the same arrow placements in the merged square demonstrate no overlap between GFP and BMAL1.

### *Bmal1* deletion in the LHb has a marginal effect on anxiety and depressive like behaviour

Tests for anxiety- and depressive-like behaviours began three weeks after surgery using the elevated plus maze test (EPM), open field test (OFT), and the sucrose preference test (SPT). Deletion of *Bmal1* in the LHb had no effect on standard measures of anxiety-like behaviours in the EPM in either males or females (Fig. 2A, B, C, & D, Table 1.). In the OFT, knockout females travel significantly less compared to controls (Fig. 2F, Table 1.), whereas male knockouts spent significantly less time in the center of the field compared to controls (Fig. 2G, Table 1.). The knockout had no effect on the latency to enter the center of the field in either males or females (Fig. 2I & J, Table 1). Lastly, in both males and females, deletion of *Bmal1* had no effect on sucrose preference, a test for anhedonia-like behaviour in mice (Fig. 2K & L, Table 1). Thus, deletion of *Bmal1* in the LHb does not significantly affect measures of anxiety and depressive like behaviours in the EPM and SPT and only marginally influences behavioural measures in the OFT, suggesting that the LHb circadian clock is not critically involved in the control of affective state.

**Table 1.** Measures of affective behaviour.

| Test | Males | Females |
| --- | --- | --- |
| **EPM: time in open arms** | $t(9) = 0.77, p = .975, n = .000$ | $t(13) = 0.92, p = .373, n = .061$ |
| **EPM: open arms entries** | $t(9) = 2.23, p = .053, n = .355$ | $t(13) = 1.35, p = .200, n = .123$ |
| **OFT: distance travelled** | $t(9) = 1.93, p = .085, n = .294$ | $t(13) = 3.35, p = .005, n = .463$ |
| **OFT: time in center** | $t(9) = 2.99, p = .015, n = .499$ | $t(13) = 1.61, p = .131, n = .166$ |
| **OFT: latency to enter** | $t(9) = 1.10, p = .299, n = .119$ | $t(13) = 1.13, p = .279, n = .089$ |
| **SPT** | $t(9) = 0.32, p = .757, n = .011$ | $t(13) = 1.075, p = .302, n = .080$ |

**Figure 2.**
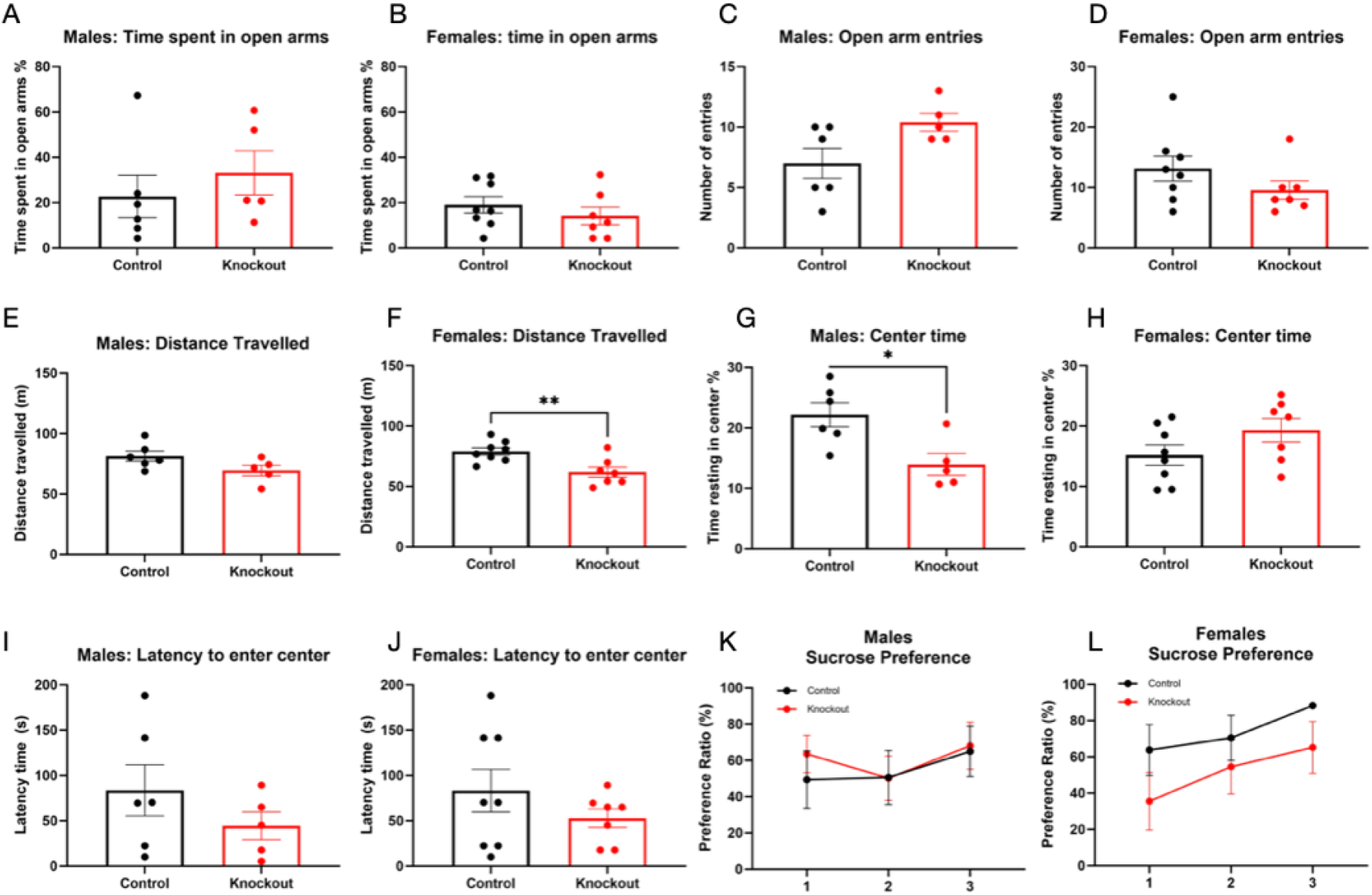
LHb *Bmal1* does not meaningfully impact affective behaviours. Males: n = 11 (5 KO); females: n = 15 (7 KO). Independent sample T-tests were run for all tests, excluding sucrose preference test. Analyses were separated by sex. (A-B) Elevated plus maze, percentage of time spent in open arms; males (A) and females (B). (C-D) Elevated plus maze, number of entries into the open arms; males (C) and females (D). (E-F) Open-field test, distance travelled; males (E) and females **p < .01 (F). (G-H) Open-field test, time resting in the center of the open-field; males *p < .05 (G) and females (H). (I-J) Open-field test, latency to enter the center of the open-field; males (I) and females (J) (K-L) Two-way ANOVA for sucrose preference across three days for males (K) and females (L).

### Sex dependent effect of *Bmal1* deletion in the habenula on alcohol consumption

The effect of *Bmal1* deletion on voluntary alcohol consumption was studied using an intermittent two-bottle choice paradigm in which mice had free access to one bottle of 15% ethanol solution and one bottle of tap water every other day for a total of 12 daily sessions. The results demonstrate sex specific effects of LHb *Bmal1* deletion on voluntary alcohol consumption. While male knockouts gradually increased their alcohol intake and preference over the 12 sessions relative to controls (Fig. 3A & C, Table 2), a marginal, statistically insignificant negative effect on consumption and preference was observed in females (Fig. 3B & D, Table 2).

**Table 2.**
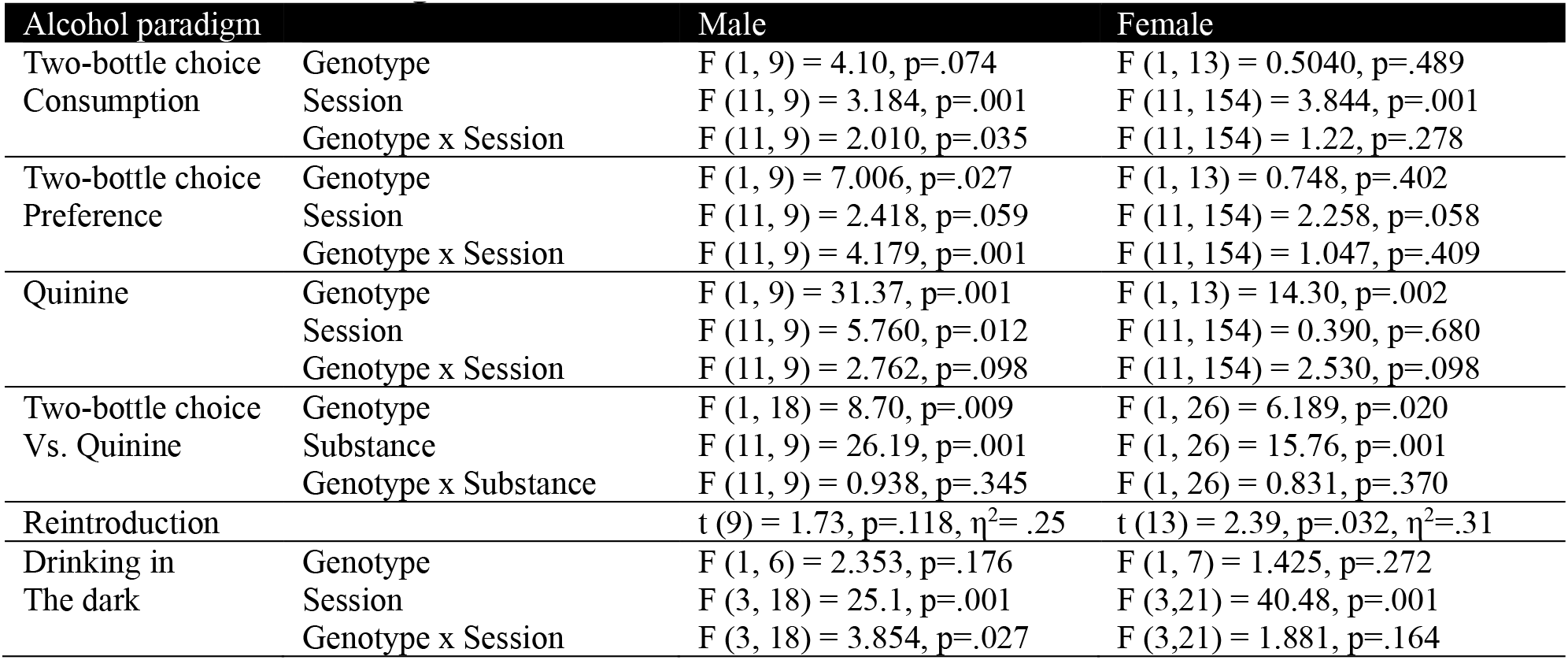
Alcohol drinking results.

**Figure 3.**
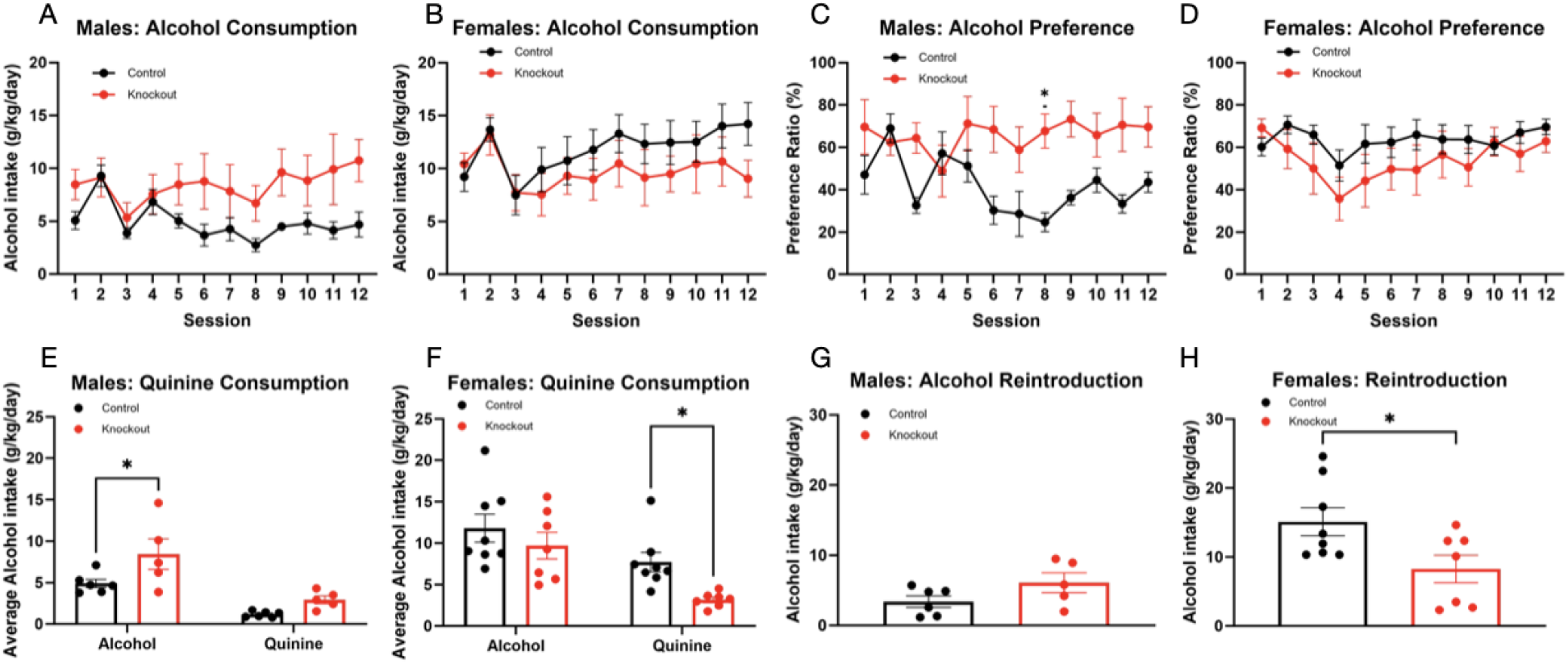
LHb *Bmal1* KO increased drinking in males, and attenuated levels in females. Males: n= 11 (5 KO); females: n= 15 (7 KO). Daily alcohol intake for males (A) and females (B) across 12 sessions of two-bottle choice paradigm. Daily alcohol preference for males (C) and females (D) calculated as ml of alcohol solution consumed/total fluid intake across 12 sessions of two-bottle choice paradigm. Alcohol intake for males (E) and females *p < .05 (F) across three sessions of two-bottle choice after the addition of quinine. Alcohol intake for males (G) and females *p < .05 (H) after a period of forced abstinence.

Two days after the last intermittent access session, all mice were given access to one bottle of 15% alcohol solution containing bitter quinine (250 uM) and one bottle of water for three consecutive days. Here, we aimed to assess if the effects of LHb *Bmal1* KO on voluntary alcohol drinking extend to the consumption of an aversive alcohol solution. The addition of quinine to the alcohol solution supressed the levels of consumption in males compared to the average levels seen in the previous 12 intermittent drinking sessions. However, despite the overall decrease in consumption, knockout males consumed significantly more of the bitter alcohol solution compared to controls (Fig. 3E, Table 2) whereas female knockouts consumed significantly less of the bitter alcohol solution relative to controls (Fig 3F, Table 2). Thus, in males, knocking out *Bmal1* in the LHb augments intake of both normal and aversive alcohol solutions. In contrast LHb knockout in females has little effect on consumption of a normal alcohol solution but represses consumption of a bitter alcohol solution.

In the last phase of this experiment, we assessed the effect of LHb *Bmal1* deletion on alcohol consumption following a period of forced abstinence, a test of relapse-like drinking behaviour. In males, both control and knockout mice consumed less alcohol during the 24-hours drinking test compared with the average levels seen across the 12 intermittent sessions, and no significant differences in intake were noted between knockouts and controls (Fig. 3G, Table 2). In contrast, knockout females consumed significantly less alcohol than controls during the 24 hours test (Fig. 3H, Table 2), pointing to a female specific inhibitory effect on drinking after forced abstinence.

### Male specific effect of *Bmal1* deletion in the habenula on alcohol binge drinking

In a separate experiment, we assessed the impact of *Bmal1* deletion in the LHb on alcohol binge drinking using the “drinking in the dark paradigm” as an experimental model (Thiele & Navarro, 2014). For this experiment, alcohol naïve knockout and control mice received a single bottle of 20% ethanol for two hours on three consecutive nights, starting two hours after lights off (ZT14-16), and for four hours on the fourth, “binge drinking” night (ZT14-18). As expected, consumption of the 20% alcohol solution increased significantly across the four sessions in all mice, regardless of sex or genotype (Fig. 4, Table 2). On the final four-hour session, male knockout mice drank nearly twice as much as controls (Fig. 4A & C, Table 2), demonstrating significant augmentation of binge drinking. In contrast, deletion of LHb *Bmal1* in females did not significantly influence alcohol consumption during the test (Fig. 4B &D, Table 2), indicating a male-specific effect of a LHb *Bmal1* KO on binge drinking.

**Figure 4.**
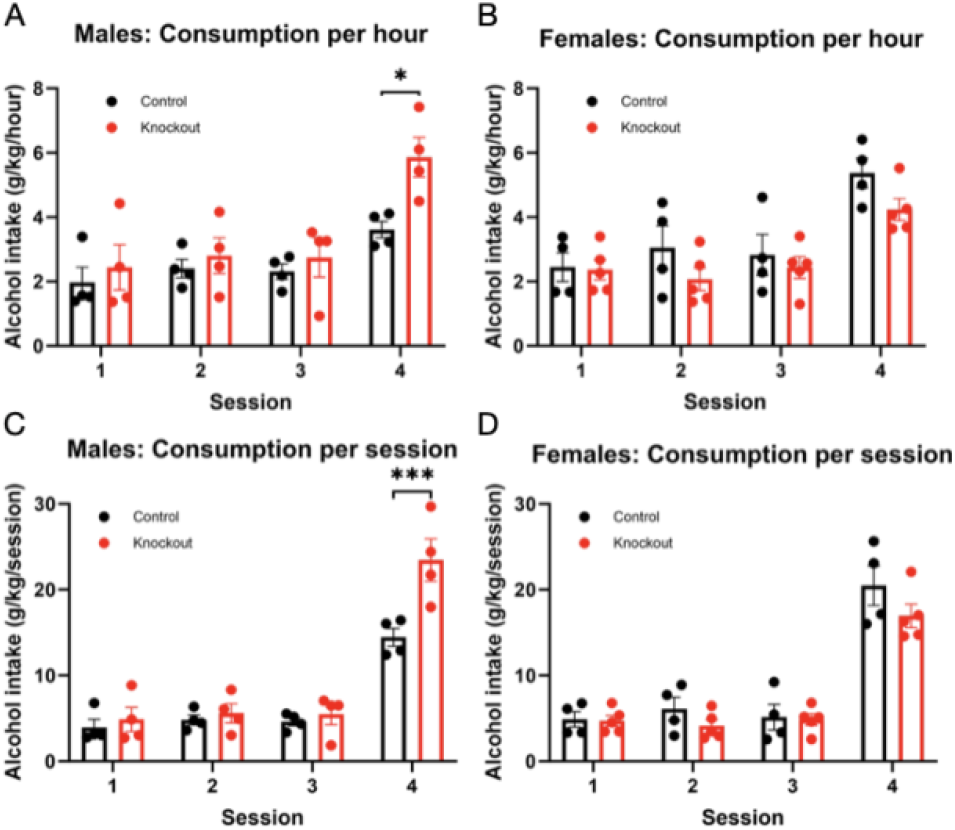
Male KO had a higher intake than controls. Males: n= 8 (4 KO); females: n= 9 (5 KO). (A) Male and (B) female “drinking in the dark” alcohol intake of control and knockout mice, normalized to hours of drinking. Per session alcohol consumption for males (C) and females (D). Sessions 1-3: 2 hours access to a 20% alcohol bottle; session 4: 4 hours access to a 20% alcohol bottle. Notes. * p<.05, *** p<.001

## DISCUSSION

This study identifies the LHb as a novel site of sex-specific effects of *Bmal1* on alcohol consumption in mice. In males, deletion of *Bmal1* in the LHb gradually augmented voluntary alcohol intake in the two-bottle choice test, attenuated the decrease in consumption of a bitter alcohol solution, and amplified binge-like alcohol consumption. In females, the same deletion did not significantly affect alcohol consumption in the two-bottle choice and binge drinking tests, but it suppressed drinking of a bitter alcohol solution and attenuated relapse-like intake following a period of forced abstinence. These results suggest that the LHb circadian clock or other *Bmal1*-dependent mechanisms normally repress alcohol consumption in males and upregulate intake under certain conditions in females. Furthermore, the findings support the hypothesis that *Bmal1*-dependent mechanisms across different brain regions contribute to sex differences in alcohol drinking in mice by downregulating intake in males and promoting heightened intake in females (de Zavalia et al., 2021).

### LHb deletion on *Bmal1* did not affect anxiety and depressive-like behaviours

Notably, the deletion of *Bmal1* in the LHb did not affect anxiety and depressive-like behaviours in the EPM and SPT, and only marginally influenced anxiety measures in the OFT. These findings mirror previous results in mice with *Bmal1* deletion in the striatum or only in the NAc (de Zavalia et al., 2021; Herrera et al., 2023; Schoettner et al., 2022), suggesting that affective state alone is not a significant determinant of the drinking phenotypes associated with *Bmal1* deletion in these regions. Interestingly, several clock genes have been associated with affective state in animal models and in humans (Benedetti et al., 2003; Chung et al., 2014; De Bundel et al., 2013; Mansour et al., 2006; Roybal et al., 2007; Russell et al., 2021; Schnell et al., 2015; Serretti et al., 2005; Zhao & Gammie, 2018), and anxiety and mood disorders commonly occur with alcohol use disorder, and the co-occurrence is associated with increased severity of both (McHugh & Weiss, 2019). Our present and previous studies suggest that any links between clock genes, affective state, and alcohol drinking behaviour are likely mediated outside the LHb-striatum circuit.

### Neural mechanisms of LHb functioning

The neural mechanisms that mediate the effect of *Bmal1* in the LHb on alcohol consumption remains to be determine. One possibility is that loss of *Bmal1* disrupts rhythmic neural communication between the LHb and the midbrain DA nuclei, altering DA synthesis and release in the striatum. Another possibility is that loss of *Bmal1*in the LHb affects the release of serotonin (5-HT) from the neurons of the raphe nuclei which, like midbrain DA neurons, have been implicated in alcohol drinking behaviour (Lovinger, 1997; Lovinger & Alvarez, 2017; Metzger et al., 2017; Sari et al., 2011). In support of the DA hypothesis, the clock in the LHb drives temporal variations in local expression of clock genes and neural activity (Guilding et al., 2010), and it plays a key role in the control of rhythmic DA synthesis and release from the neurons in the VTA and SN (Ji & Shepard, 2007; Langel et al., 2018; Matsumoto & Hikosaka, 2007). Release of DA from these striatum-projecting neurons has been implicated in the circadian control of striatal clock gene expression and in the regulation of alcohol drinking behaviour (Korshunov et al., 2017; McClung, 2007; Parekh et al., 2015; Tang et al., 2022; Verwey et al., 2016; Hood et al, 2010; Gravotta et al., 2011). Previous work indicates that a *Bmal1* knockout in both the whole striatum and the NAc specifically altered drinking behaviour (de Zavalia et al., 2021; Herrera et al., 2023). Thus, a molecular circadian clock in the LHb has the potential to indirectly influence striatal DA signaling, striatal clock gene expression, and striatum-mediated alcohol consumption.

### Female mice exhibited habitual alcohol drinking

To our knowledge, this is the first evidence of LHb/Bmal-mediated sex differences in alcohol consumption. However, LHb-dependent sexual dimorphisms have been found in various neural and behavioral functions of the LHb. For example, there are sex differences in the inhibitory influence of the LHb on the midbrain DA neurons (Bell et al., 2023), and in LHb influence on social communication (Rigney et al., 2020), parental behaviour (Lecca et al., 2014), and stress responses (Zhang et al., 2018). As such, the LHb clearly influences certain neural and behavioural processes in a sex-dependent manner.

Here, female control mice retained high alcohol consumption throughout all paradigms, including the addition of quinine and after forced abstinence. This indicates that lowering the incentive value of the reward (i.e. alcohol) did not affect intake in control females. In animal models, persistent drug taking despite aversion or changes in reward value is defined as habitual drinking, a model of addiction-like behaviour (Dickinson, 1985; Vena et al., 2020). This shift to a habitual behaviour is associated with a change from ventral (nucleus accumbens; NAc) to dorsal striatal control of drug taking (Balleine & O’Doherty, 2010; Chen et al., 2011; Graybiel & Grafton, 2015; Smith & Graybiel, 2016). This indicates that in females, *Bmal1* in the LHb may act by supressing the shift from ventral to dorsal striatal control, inhibiting the shift to addiction-like drinking. While KO males demonstrated increased intake and preference after repeated alcohol exposure, they were significantly detered by a change in the reward value (quinine and forced abstinence). As such, males do not meet the criteria for addiction-like alcohol drinking. Thus, we propose that *Bmal1* in the LHb of male mice may typically supresses NAc related alcohol drinking behaviour but does not impact addictive-like drinking.

Notably, the difference in KO and control females was increased after the addition of quinine and after forced abstinence, the aversive paradigms. As the LHb plays an important role in aversive conditioning and acts as the so-called “negative reward center”, this difference is likely due to a sex-specific role of *Bmal1* in mediating LHb functioning (Fu et al., 2017; Mondoloni et al., 2022; Zuo et al., 2014). The LHb typically has a weaker inhibitory influence on midbrain DA functioning in females (Bell et al., 2023) and estrogen in the LHb mediates the response to stressful stimuli (Calvigioni et al., 2023). *Bmal1* may mediate LHb functioning through its control over estrogen. *Bmal1* binds to the E-Box in the promoter region of estrogen receptor β, controlling rhythmic oscillations (Cai et al., 2008). As estrogen receptors are denser in the female LHb (Bell et al., 2023), *Bmal1* may alter LHb functioning in a sex-dependent manner, thereby contributing to the observed sexually dimorphic drinking behaviour.

### D2 receptors play a role in increased alcohol drinking behaviour

Sex differences in DA functioning have been well-established, especially in the striatum (Becker, 1990; Calipari et al., 2017; Dewing et al., 2006; McArthur et al., 2007; Walker et al., 2006; Zachry et al., 2021). Previously, we found that deleting *Bmal1* in the LHb indirectly alters striatal DA functioning (Goldfarb et al., *in prep*). Striatal DA is essential for clock gene functioning (Hood et al., 2010; Imbesi et al., 2009) and striatal clock genes including *Bmal1* and *Per2* impacts alcohol drinking behaviour in a sexually dimorphic manner (de Zavalia et al., 2021). As such, the sexually dimorphic drinking patterns found here may in part be due to LHb *Bmal1* mediated changes in DAergic functioning.

DA in the striatum functions primarily through two main pathways, the direct D1 pathway and indirect D2 pathway. The interaction between DA and clock genes in the striatum is also cell type specific, as D1 and D2 receptor agonists influence clock gene functioning differently (Hood et al., 2010; Imbesi et al., 2009). Moreover, stimulation of D1 and D2 receptors differentially impact alcohol drinking behaviour, emphasizing the distinct role of these circuits (Cheng et al., 2017). Therefore, the impact of LHb *Bmal1* on alcohol drinking behaviour may be through cell type specific DAergic mechanisms in the striatum. For example, D2 receptors on striatal medium spiny neurons (MSNs) have been linked to alcohol use disorder in humans (Hietala et al., 1994; Tupala et al., 2001; Volkow et al., 2019). Likewise, chronic alcohol consumption in mice reduced D2 receptor availability (Feltmann et al., 2018), and overexpression of D2 receptors in rats led to reduced alcohol preference and intake (Thanos et al., 2001). After selective deletion of D2 receptors on MSNs, both male and female mice increased alcohol preference (Bocarsly et al., 2019). This is theorized to be due to the subsequent imbalance between the direct and indirect (D1 and D2) pathways, resulting in altered striatal circuitry and increased vulnerability to alcohol. The present findings demonstrate that *Bmal1* in the LHb typically decreases the propensity for alcohol drinking in males and increases it in females. This, along with our previous findings that LHb *Bmal1* influences DA functioning, allows us to speculate that *Bmal1* in the LHb may impact alcohol drinking behaviour by indirectly altering the balance of striatal DA circuitry.

The complexity of these interactions highlights the need for a more comprehensive understanding of the molecular and neural circuit mechanisms underlying the sex-specific regulation of alcohol consumption by *Bmal1*. These insights are essential for developing targeted therapeutic strategies for alcohol use disorders, which often exhibit sex differences in prevalence and severity. By elucidating the specific pathways and molecular players involved, future research can pave the way for more effective treatment.

## MATERIALS AND METHODS

### Subjects

*Bmal1* floxed mice (B6.129S4(Cg)-Arntl^tm1Weit^/J) were originally obtained from Jackson Laboratories. 43 mice were used in total, 19 males and 24 females. Animals were divided into two groups, A and B. Group A consisted of 11 males (6 control/ 5 knockout) and 15 females (8 control/ 7 knockout). Group B consisted of 8 males (4 control/ 4 knockout) and 9 females (4 control/ 5 knockout).

Male and Female mice were group-housed under a 12:12 light-dark (LD) cycle at 21 ± 2°C and 60% relative humidity with food and water available *ad libitum* prior to the stereotactic delivery of viral vectors at 12 – 16 weeks of age. The bedding of the cages was changed every week. Animals were housed individually after the surgery under the same housing conditions as described above.

All experiments were conducted according to the guidelines and requirements of the Canadian Council on Animal Care (CCAC) and approved by the Concordia University ethics committee (AREC number 3000256).

### Stereotaxic Surgeries

*Bmal1* floxed mice were anesthetized by intraperitoneal injection of Ketamine (100 mg/kg body weight) and Xylazine (10 mg/kg body weight) solution and received subcutaneous injections of Ketoprofen (5 mg/kg body weight) as post-operational analgesia. Animals were placed in a stereotaxic apparatus (KOPF, Tujunga, CA, USA) and received bilateral intra-LHb microinjections of recombinant viruses (AP: -1.65, ML: 0.4, and DV: -3.1) using a 30-gauge needle attached to a 10 ul Hamilton syringe connected to a micropump (Harvard Aparatus). The injector was inserted at a 0° angle and the virus was delivered at a rate of 100 nl/min (120 nl total volume). For the generation of conditional *Bmal1* knockout, mice were given recombinant viral vectors expressing Cre and enhanced green fluorescence protein (eGFP) (AAV2/9-CAG-Cre-eGFP, 1×10^12^ vg/ml). Control animals received intra-LHb viruses expressing eGFP only (AAV2/9-CAG-eGFP, 1×10^12^ vg/ml). The injector was left in place for five minutes following the injection to optimize diffusion. Mice were allowed to recover for three weeks before behavioural testing. LHb-specific expression of viral vectors was evaluated at the end of the experiment. Animals with missing, incomplete, or off-region eGFP expression were excluded from all experiments.

### Behavioural procedures, Ethanol consumption and Sucrose preference

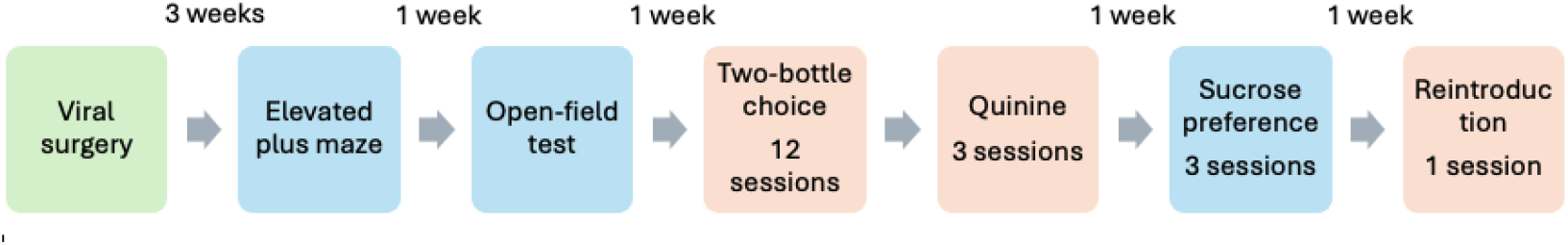

Group A (11 males (5 KO); 15 females (7 KO) were assessed on the elevated plus maze (EPM), open-field test (OFT), two-bottle choice, quinine, sucrose preference test, and reintroduction session. Animals were euthanized after the reintroduction session and brains were collected for viral validation.

Alcohol naïve group B (8 males (4 KO) 9 females (5 KO)) was assessed only on the drinking in the dark paradigm. Animals were euthanized after the fourth session and brains were collected for viral validation.

### Elevated Plus Maze

The elevated plus maze was used as an assay for anxiety-like behaviours. The “+”-shaped maze was positioned 40 cm above the ground with two enclosed arms (6 × 29.5 cm), two open arms (6 × 29.5cm) and a center area (6 × 6 cm). White methacrylate floors and black walls (15 cm in height) lined the bottom and enclosed arms of the maze, respectively. The test is based on mice’s natural aversion to heights and open areas (Walf and Frye, 2007). Mice were habituated to the testing room by leaving them in their cage for an hour before the start of the experiment.

Then, each mouse was placed in the center area of the elevated plus maze (Harvard Apparatus, Holliston, MA, USA) facing the closed arm and allowed to explore the maze for 5 minutes while video recorded (Samsung Galaxy A5 2017 phone). Video files were analyzed as described in Schoettner et al. (2022). Time spent in the open arms and number of entries into the open arms were assessed. The test was conducted at Zeitgeber time 2 (ZT2, ZT0 represents the time of lights-on).

### Open field test

The open field test was used to assess anxiety-like behaviour and motor activity. After a 1-hour habituation period in the experimental room, mice were placed in the corner of the open field arena (45 × 45 × 60 cm, Panlab, Barcelona, Spain) facing the wall. The open field was equipped with infrared beams to track horizontal activity over 30 minutes using the ACTItrack software (Panlab, Barcelona, Spain). At the end of the session, animals were weighed and returned to home cages while the arena was wiped with a 70% ethanol solution (v/v in tap water) to remove any residues and olfactory cues before the next set of animals were tested. Total distance travelled, permanence time in the center of the open field, and the latency to enter the center were assessed. The test was conducted at ZT 2.

### Two bottle choice

Voluntary alcohol consumption was studied using the intermittent two-bottle choice test as described previously (de Zavalia et al., 2021). Briefly, mice were habituated to drinking water from two bottles for three days, one week after the last behavioural test. They were then given access to one bottle of alcohol solution (15% v/v, in tap water) and one bottle containing tap water every other day, for 12 sessions. The position of the alcohol and water bottles was altered each session to control for potential side preference. All mice were given access to two water bottles during the alcohol-off days. Alcohol and water intake (g/kg body weight/day) and alcohol preference (ml of alcohol solution consumed/total fluid intake) were measured daily and calculated at the end of the 12 sessions. Body weights were collected weekly during cage changes.

### Quinine and alcohol

Continuous drinking despite aversive consequences is a DSM-5 diagnostic criteria for alcohol use disorder. One way this is modeled in animal voluntary drinking paradigms is through the addition of quinine to the alcohol solution. Quinine is a bitter tasting substance and its addition to ethanol alters the valence of alcohol, making the solution less palatable. The same procedure as described above for two bottle choice was used. In brief, mice were given one bottle of 15% ethanol containing 250 uM quinine in tap water (v/v) and one water bottle for three consecutive days. Bottle placement was swapped every day. Alcohol and water intake (g/kg body weight/day) were measured daily and calculated at the end of the three sessions. Body weights were collected in morning before the first session.

### Sucrose preference

A sucrose preference test was performed one week following the alcohol/quinine test. To minimize the effects of neophobia as a confounding factor, animals were given two bottles of 1% sucrose solution (v/v in tap water) for 4 hours the day before testing, and the volume of consumed sucrose solution was assessed to ensure all animals were drinking. During the test, mice were given unlimited access for 24-hours to one bottle of tap water and one bottle of 1% sucrose solution for three consecutive days, with the position of the sucrose and water bottles switched every day to control for side preference. The ratio of the sucrose solution consumed relative to the total fluid intake was used as a measure of sucrose preference (V_sucrose solution_/(V_sucrose solution_ + V_water_)).

### Reintroduction

After a period of forced abstinence (1 week after SPT; 10 days after the completion of the 3-day quinine/ alcohol paradigm) mice were reintroduced to one bottle of 15% alcohol in tap water (v/v) and one bottle of tap water for a single 24-hour session to investigate the effect of LHb *Bmal1* deletion on relapse-like drinking. Alcohol and water intake (g/kg) were measured and calculated at the end of the session. Body weights were collected after the session.

### Drinking in the dark test

To assess binge drinking behaviour, alcohol naïve mice underwent a four-day drinking in the dark paradigm (Thiele et al., 2014). In brief, mice were given 2h access to one bottle of 20% ethanol in tap water (v/v) for 3 consecutive days, starting 2 hours into the dark phase (ZT14), and for 4 hours on the fourth day. Body weights were measured before the first session and at the end of the last session. Alcohol intake (g/kg body weight/day) was measured each day and calculated at the end of the final session.

### Immunofluorescence

Animals were deeply anesthetized by exposure to an atmosphere of isoflurane and perfused transcardially by cold saline (0.9% sodium chloride, pH 7.2) followed by paraformaldehyde solution (PFA, 4% in 0.1M phosphate buffer, pH 7.2) using an infusion pump. Brains were then dissected and postfixed in PFA for 22 – 24h at 4°C. 30 μm coronal section of brain tissue were collected using a Leica vibratome and analyzed under a fluorescent microscope to validate region-specific viral vector delivery. Brain slices were stored at −20 °C in Watson’s cryoprotectant (Watson et al., 1986) for immunofluorescence imaging.

For this, free-floating sections previously kept in Watson’s cryoprotectant were rinsed for 10 minutes in phosphate buffered saline (PBS, pH 7.4). Following this, sections were washed three times in PBS containing 0.3 % Triton-X (PBS-Tx) for 10 min, followed by an incubation in blocking solution (3 % milk powder, 6 % normal donkey serum (NDS) in PBS-Tx) for one hour at room temperature with mild agitation. Primary antibody solution was prepared by diluting the primary antibody for BMAL1 (rabbit anti-BMAL1, 1:500, NB100-2288, Novus Biologicals) in PBS-Tx containing 3 % milk powder and 2 % NDS. Brain sections were incubated for 1.5 hours at room temperature. Following three washes in PBS-Tx, brain sections were incubated at room temperature in secondary antibody solution containing donkey anti-rabbit IgG Alexa Flour 647 (1:500, Thermo Scientific™)) diluted in PBS-TX (3 % milk powder and 2 % NDS) for one hour. Brain sections were mounted on gel coated microscope slides and cover slipped using a mounting media containing DAPI (ProLong™ Diamond Antifade Mountant, Thermo ScientificTM) after three washes 10-min in PBS-Tx and a final wash in PBS for 10 min. Brain sections were stored at 4 °C until imaging under an Olympus 10FI confocal microscope. A ratio between BMAL1 expressing cells over GFP expressing stained cells was used to calculate rate of viral infection. Representative images were prepared using Image J.

### Data analysis and statistics

The collected raw data from the behavioural tests and alcohol consumption experiments were further processed and analyzed using Prism 9 (GraphPad Software, San Diego, CA, USA). Unpaired two-tailed t tests and repeated measures ANOVAs were used to compare differences between groups. Significant main effects and interactions of the ANOVAs were further investigated by the appropriate post-hoc test. The level of statistical significance was set at P < 0.05. Exact P values and further details are given in the tables above. The outcomes were depicted as mean ± standard error of the mean (SEM).

## Acknowledgements

This work was funded by grants from the Canadian Institutes of Health Research (S.A). All microscopy was performed at the Concordia Centre for Microscopy and Cellular Imaging at Concordia University, Montreal (special thanks to Dr. Chris Law)

## Author contributions

K.S., S.A., and C.G. conceived and designed the study; C.G., V.H., and A.M. performed the experiments; C.G., K.S., and S.A. analyzed and interpreted the data, and wrote the manuscript; S.A. and K.S. Supervision.

